# Epigenetic CpG Duplex Marks Probed by an Evolved DNA Reader via a Well-Tempered Conformational Plasticity

**DOI:** 10.1101/2022.10.06.511192

**Authors:** Himanshu Singh, Chandan K. Das, Benjamin C. Buchmuller, Simone Eppmann, Lars V. Schäfer, Daniel Summerer, Rasmus Linser

## Abstract

5-methylcytosine (mC) and its TET-oxidized derivatives exist in CpG dyads of mammalian DNA and regulate cell fate, but how their individual combinations in the two strands of a CpG act as distinct regulatory signals is poorly understood. Readers that selectively recognize such novel “CpG duplex marks” could be versatile tools for studying their biological functions, but their design represents an unprecedented selectivity challenge. By mutational studies, NMR relaxation, and MD simulations, we here show that the selectivity of the first designer reader for an oxidized CpG duplex mark hinges on precisely tempered conformational plasticity of the scaffold adopted during directed evolution. Our observations reveal the critical aspect of defined motional features in this novel reader for affinity and specificity in the DNA/protein interaction, providing unexpected prospects for further design progress in this novel area of DNA recognition.

## Introduction

Cellular differentiation to stable, tissue-specific phenotypes despite identical genetic material is a prerequisite for the development of multicellular organisms. This is achieved by coordinated gene expression regulation via chromatin modification, such as the epigenetic modification of DNA nucleobases. In mammals, 5-methylation of cytosine by DNA methyltransferases (DNMTs) plays essential roles in differentiation, development, X-chromosome inactivation, and genomic imprinting; consequently, aberrant DNA methylation has been linked to multiple diseases, including cancer.^1–2^ Enzymatic oxidation of mC (Fig. 1A) to 5-hydroxymethylcytosine (hmC), 5-formylcytosine (fC), and 5-carboxycytosine (caC) is catalyzed by Ten-Eleven-Translocation (TET) dioxygenases and results in particularly high levels of oxidized mCs in embryonic stem cells and the brain.^3^ These oxidized mC derivatives have been shown to exert regulatory functions in multiple contexts.^4–6^ Mammalian cytosine modification by DNMTs and TETs occurs predominantly in palindromic CpG dyads and can theoretically give rise to 15 different symmetric and asymmetric combinations of cytosine 5-modifications across the two CpG strands.^7^ However, despite the established general roles of oxidized mCs as chromatin regulators, it is poorly understood how their individual combinations in CpGs act as distinct regulatory signals, for example, by differentially modulating interactions with the large number of double-stranded DNA-binding chromatin proteins. Reader proteins that selectively interact with novel CpG duplex marks could serve as fundamental tools for studying their biological functions,^8–11^ but their design opens a new aspect in the field of DNA recognition that poses formidable selectivity challenges. We have recently reported the first designer reader for such a TET-associated CpG duplex mark. This protein has been evolved from a methyl-CpG-binding-domain (MBD)^12^ and selectively recognizes the asymmetric combination hmC/mC in the context of all fifteen possible CpG duplex marks.^13^

**Fig. 1:**
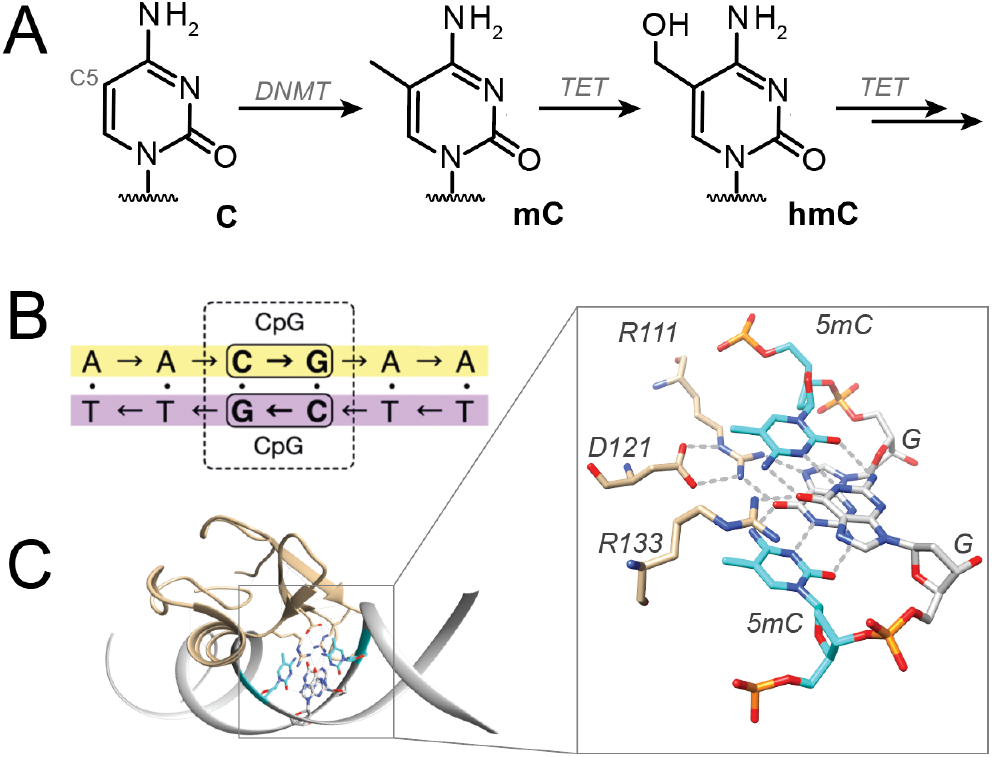
Natural selection of mC/mC DNA by methyl-CpG-binding domains (MBDs). **A)** Cytosine modifications at CpG dyads, generated via DNMT and TET enzymes. **B)** Depiction of a CpG dyad, in which cytosine can be epigenetically modified either symmetrically or asymmetrically. **C)** Recognition of mC/mC dyads by MBDs via two H-bonds each from Arg111 and Arg133 to guanosine.^14^

The “core” MBD family proteins share a conserved domain of 70-80 residues and include the proteins MBD1–4 and Methyl-CpG-binding protein 2 (MeCP2).^12^ The latter represents a largely disordered DNA-binding protein for which loss-of-function mutations are associated with the neurological developmental disease Rett syndrome (RTT).^15–16^ Its high-affinity interaction with mC/mC DNA hinges on two Arg fingers that both form two H-bonds to the guanosine of the CpG (Fig. 1C).^14^ Previous work on MBDs has shown that the stability of the three-dimensional fold is exceptionally susceptible to simple point mutations in the center of the hydrophobic core.^17^ Now, directed evolution of MBDs has recently been suggested as a viable path to generating reader proteins specifically targeting previously inaccessible combinations of epigenetic DNA modifications in CpG dyads, with the prospect of providing a platform for their genome-wide identification and mapping.^13, 18^

Directed evolution experiments for selecting hmC/mC readers from an MeCP2 mutant library revealed the replacement of a hydrophobic core residue (Val122) with Ala as a critical mutation, emerging in addition to a modified DNA binding interface (K109T/S134N, Fig. 2A).^13^ The V122A mutation in the TAN triple mutant established high hmC/mC DNA-binding affinity (~10 nM) and specificity in electromobility shift assays (EMSA). In contrast, a second selected mutant (TCN) exhibited more promiscuous binding of both mC/mC and hmC/mC, even though it differed only by one nonpolar Cys residue at a core position that does not interact with DNA in the wt protein. This surprising role of a core residue in designed CpG duplex readers as potent determinant for selective target recognition reveals a fundamental lack of molecular-level understanding and unravels a considerable pitfall of structure-based approaches for the design of this novel class of epigenetic reader proteins.

**Fig. 2:**
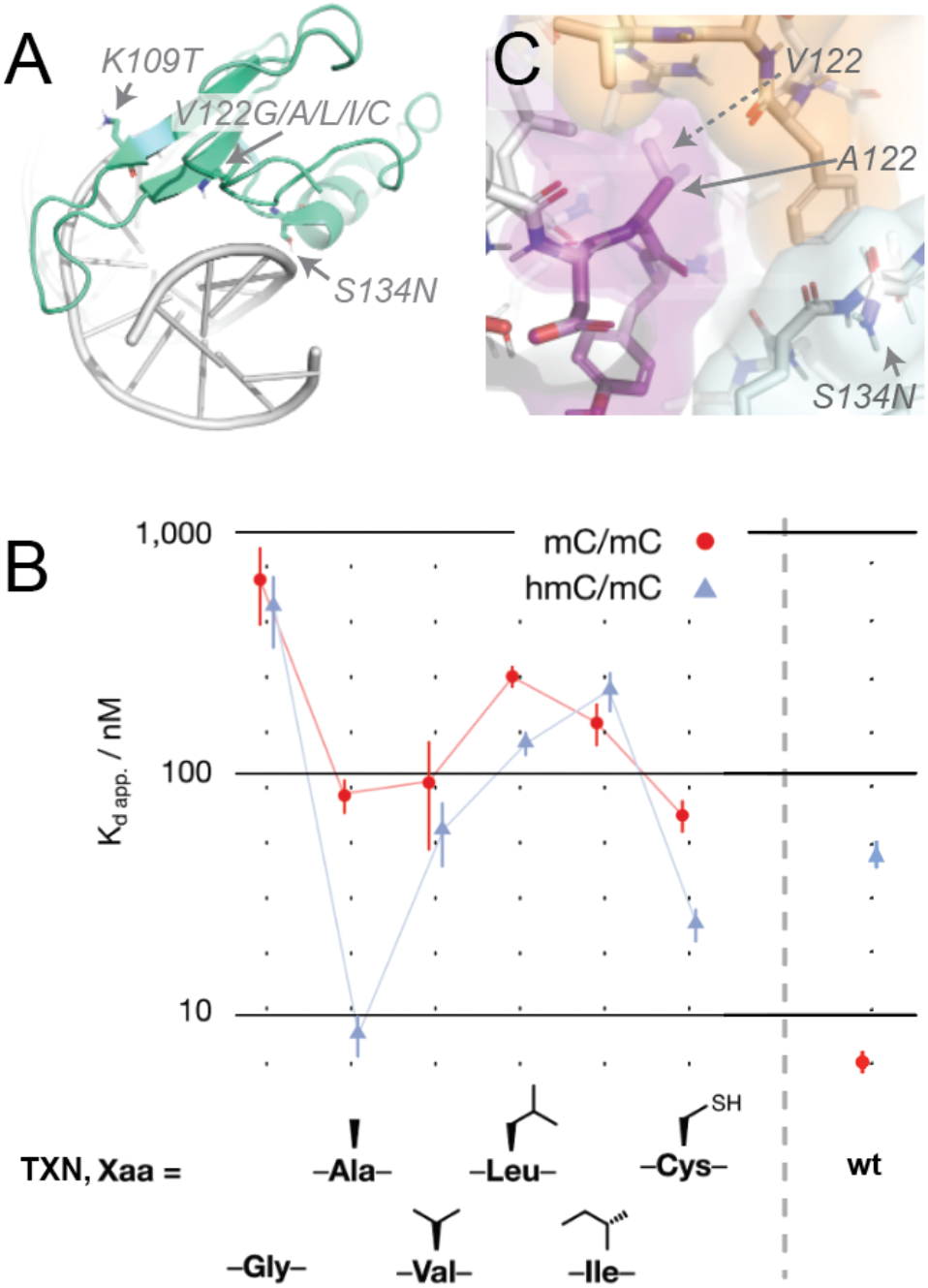
Engineering of hmC/mC selectivity into the natural MeCP2 reader. **A)** Mutation sites in wt MBD (pdb: 3c2i). **B)** Affinity for different triple mutations (K109T/V122X/S134N, “TXN”) in comparison to wt MBD (right), determined from EMSA shift essays. (X = Ala, second column, represents “TAN”; X = Val, third column (“TVN”), represents *double* mutation of the wild type.) **C)** Position 122 (the spatially more demanding Val shown in semitransparent) defines the central inter-subunit interface between extended sheet (purple, res. 104 – 126), helix 1 (green, 132 – 146), and C-terminal domain (orange, 148 – 161).

## Results

To study the peculiar influence of hydrophobic-core residues on the functional level, we generated mutants with different smaller (Gly, Ala) or larger (Val, Ile, Leu) residues at position 122, maintaining mutations required in the DNA binding interface. We expressed the respective MBDs (residues 87-190) from human MeCP2 and measured dissociation constants (K_D_) for mC/mC and hmC/mC-containing dsDNA by electromobility shift assays (EMSA, Figs. 2B and S1). Only the selected TAN mutant exhibited a high affinity (8 ±2 nM) and selectivity for its new, hmC/mC-containing target, whereas the TCN mutant exhibited a slightly lower selectivity. Strikingly, any deviation from this narrow steric space led to a dramatic loss of selectivity, most pronounced for the TGN mutant. Importantly, both MBDs with the wild type residue Val at position 122 (TVN and wt KVS) exhibited equally poor (~100 nM) binding of hmC/mC, but differed drastically in the binding to the canonical wild type target mC/mC, which was not bound by TVN anymore. Strikingly, this clear requirement for a specific steric demand at position 122 was observed despite the fact that it lies in a secluded element not interacting with DNA (Fig. 2C).

To explore the molecular underpinnings of the modulating role of the protein core for selectivity of its interface, we conducted NMR and MD simulation studies. Wild-type MBD, the hmC/mC reader TAN, as well as the TVN mutant were overexpressed in doubly-labeled ^15^N, ^13^C minimal media and purified using affinity and size-exclusion chromatography. Backbone and sidechain ^1^H, ^13^C, and ^15^N resonance assignments were obtained for wt, TAN and TVN mutants, as well as the DNA-bound form of TAN via a series of triple-resonance NMR experiments in solution (see the SI). The assigned ^15^N-^1^H HSQC spectra of apo wt and TAN triple mutant are shown in Fig. S2. Resonance assignments have been deposited for the wt, TVN, and TAN into the BMRB under accession codes 51548, 51547, and 51020, respectively.

### Ground state structure of the designed hmC/mC reader

The apo structure of the artificial hmC/mC reader TAN was determined by NMR spectroscopy using 698 internuclear distances from ^15^N-edited and aliphatic or aromatic ^13^C-edited NOESY spectra and 108 torsion angles. Backbone and hydrophobic side chains are well defined both in the helical domain and in the extended β-sheet, except for a loop region (residues 111–119, termed loop L1 in the following, compare Fig. S3) and N- and C terminal residues (Fig. 3A). Whereas the secondary-structural elements of the ordered regions in the hmC/mC reader TAN (compare Fig. S4) are identical to the wt, the relative orientation of the helical domain to the extended β-sheet remained ambiguous: Residual dipolar couplings (^1^D^NH^), obtained from partial alignment via Pf1 bacteriophages and evaluated using singular-value decomposition as implemented in PALES^19^ (see details in the SI), are mainly in agreement with the wt crystal structure (PDB 3c2i). However, significant deviations are observed for residues 134, 140, 144, and 145 in helix a1 (Figs. 3B/C), even though RDCs for residues 138, 142, and 143 on the back of the helix are as expected. Including all RDCs into structure calculation yields poor conversion, which agrees with an ambiguous orientation of helix α1 in a dynamic ensemble (see below). All of this is in contrast to the double mutant TVN, which shows RDCs exactly matching the values expected for the wt mC/mC reader (Fig. 3B). (An assigned HSQC spectrum of TVN is part of Fig. S12.) Fig. 3A depicts the backbone structures of the 20 lowest-energy conformations with a precision (root mean square deviation, RMSD) of 0.7 Å for the backbone and 1.4 Å for all heavy atoms (including residues 102–154 apart from loop L1), deposited into the PDB under accession code 8AJR. This applies to calculations including all RDCs except for the helical domain (compare Fig S5). Even though in this case the *R^2^* to the wt structure is 0.92, the precise inter-domain orientation remains undetermined (center panel of Fig. 3A). The statistics for the 20 final water-refined structures are shown in Table S1. Finally, the average structures observed in MD simulations (See more details for the MD studies below.) of apo proteins (in the absence of DNA, see Fig. 3D, also compare Fig. S17B) also point to slight deviations in the tertiary structure of the reader, the relative arrangement of the helix being slightly altered.

**Fig. 3:**
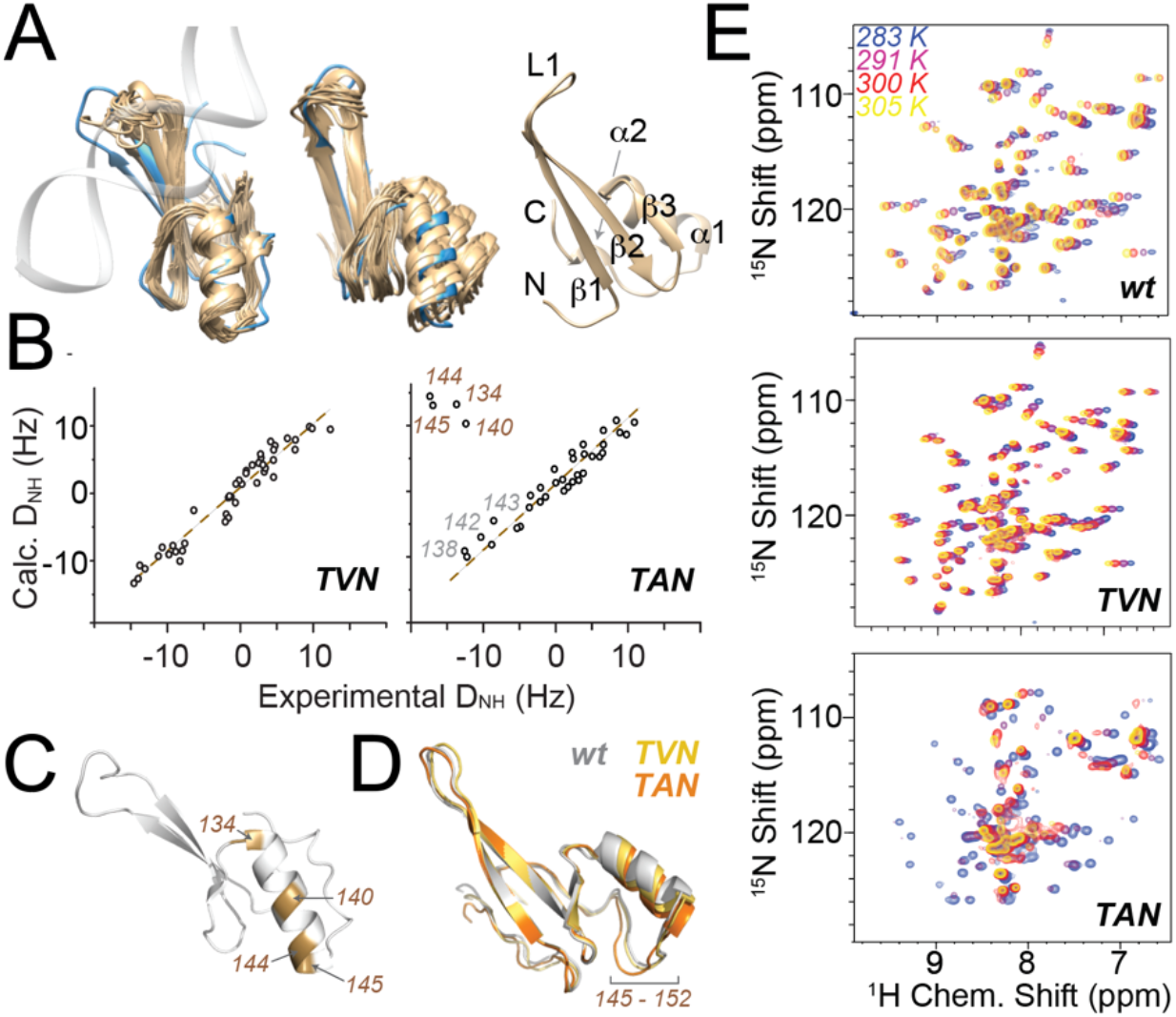
Assessment of structural properties of the apo hmC/mC reader. **A)** Solution NMR structural ensemble of the hmC/mC reader TAN, showing the 20 minimal-energy structures aligned either over all residues (left, crystallographic DNA-bound structure 3c2i in blue) or with respect to the β-sheet (center), and the lowest-energy NMR structure with secondary-structural annotations (right, slightly different angle). **B)** Residual dipolar couplings for double mutant TVN (left) and triple mutant TAN (right) correlated with theoretical values derived for the structure of the wild-type reader. **C)** Residues with deviating RDCs depicted on 3c2i. **D)** Average MD structures of the different apo proteins over 2.5 μs, highlighting structural deviations in the helix and subsequent residues (also see Fig. S17). **E)** Temperature dependence of wt (top), TVN (center), and TAN HSQC spectra (bottom), suggesting strong conformational exchange for TAN but not for wt and TVN. Structures in C and D from slightly different perspectives.

To our surprise, at a temperature of 20°C or above, many protein resonances of the hmC/mC reader TAN reversibly broaden beyond detection, indicating the presence of alternate conformational states on the μs-to-ms timescale (Fig. 3E). By contrast, no significant indication of temperature-dependent exchange broadening is observed over a wide range of temperatures in the wt protein or TVN, which carries only the mutations directly involved in the DNA interactions, rendering its binding affinity to hmC/mC almost 10-fold lower than in TAN (Figs. 2B and S1). To assess the details of conformational rearrangements in the high-affinity TAN reader, we closely examined motion occurring on the ps-ns time scale as well as in the μs-ms regime, employing a large array of ^15^N relaxation, relaxation dispersion (RD), and chemical-exchange saturation transfer (CEST) data.

### The apo hmC/mC reader accesses alternate conformational states

[^15^N, ^1^H] heteronuclear NOE, ^15^N *R*_1_, and ^15^N *R*_2_ relaxation for TAN and wt MBD are shown in Fig. S3 and S4, respectively. The distribution of hetNOE and *R*_1_ rates, reporting on ps-ns timescale dynamics, confirm the architecture of the domain with respect to its expected mobile N-terminus, extended C-terminus, and loop L1. More interestingly, transverse relaxation (*R*_2_) is, in addition, sensitive to slower motions and reflects conformational-exchange processes on the μs-ms timescale. Whereas *R*_2_ rates in wt simply mirror the fast-timescale mobility observed in *R*_1_ and hetNOE, significant deviations in TAN reveal robust conformational exchange throughout the sequence. To assess the timescale of motion for the exchange, we carried out ^15^N constant-time CPMG relaxation dispersion experiments. A dispersive nature in the dispersion profiles is the signature of conformational exchange on the μs–ms timescale between states with different chemical shifts. For wt, a small number of residues show the incidence of modest conformational exchange (Figs. 4A and S6), with global RD on a timescale of around 200 μs (*k_ex_* of 5207 ± 356 s^-1^). In the hmC/mC reader TAN, strikingly, these exchange contributions are fourfold slower and more excessive than in the wild-type – with a timescale of about 800 μs (*k_ex_* 1240 ± 10 s^-1^) at 18 °C, higher total *R*_2_ rates up to 60 s^-1^, and substantial exchange contributions *R*_ex_ up to >40 s^-1^ – widely surrounding the structural elements in loop L1, β1, β2, and β3 strands, and a1 residues. Figs. 4A and B display *R*_ex_ mapped onto the structure and exemplary dispersion profiles (selecting the three mutation sites), respectively; Fig. S7 provides further dispersion profiles for TAN. RD data were fitted individually, assuming either a two-site exchange model or the absence of exchange, dependent on the corrected Akaike information criterion. See Fig. S8 for an overview about the residues with significant exchange contributions in both wt and TAN. Interestingly, the only difference between the constructs is V122 vs. A122, which mutation hence allows a strong increase in conformational exchange. Neither V122 nor A122 backbone sites show dispersion themselves, reflecting the role of the side chain as a lever for the dynamics, the amide not being exposed to differential chemical environments itself (Fig. 4A/B center). Finally, the exchange dynamics ceases upon DNA binding. (See details regarding complex formation below.) Apparently, in the apo protein, an exchange occurs between different conformations, of which only one is relevant within the complex.

**Fig. 4:**
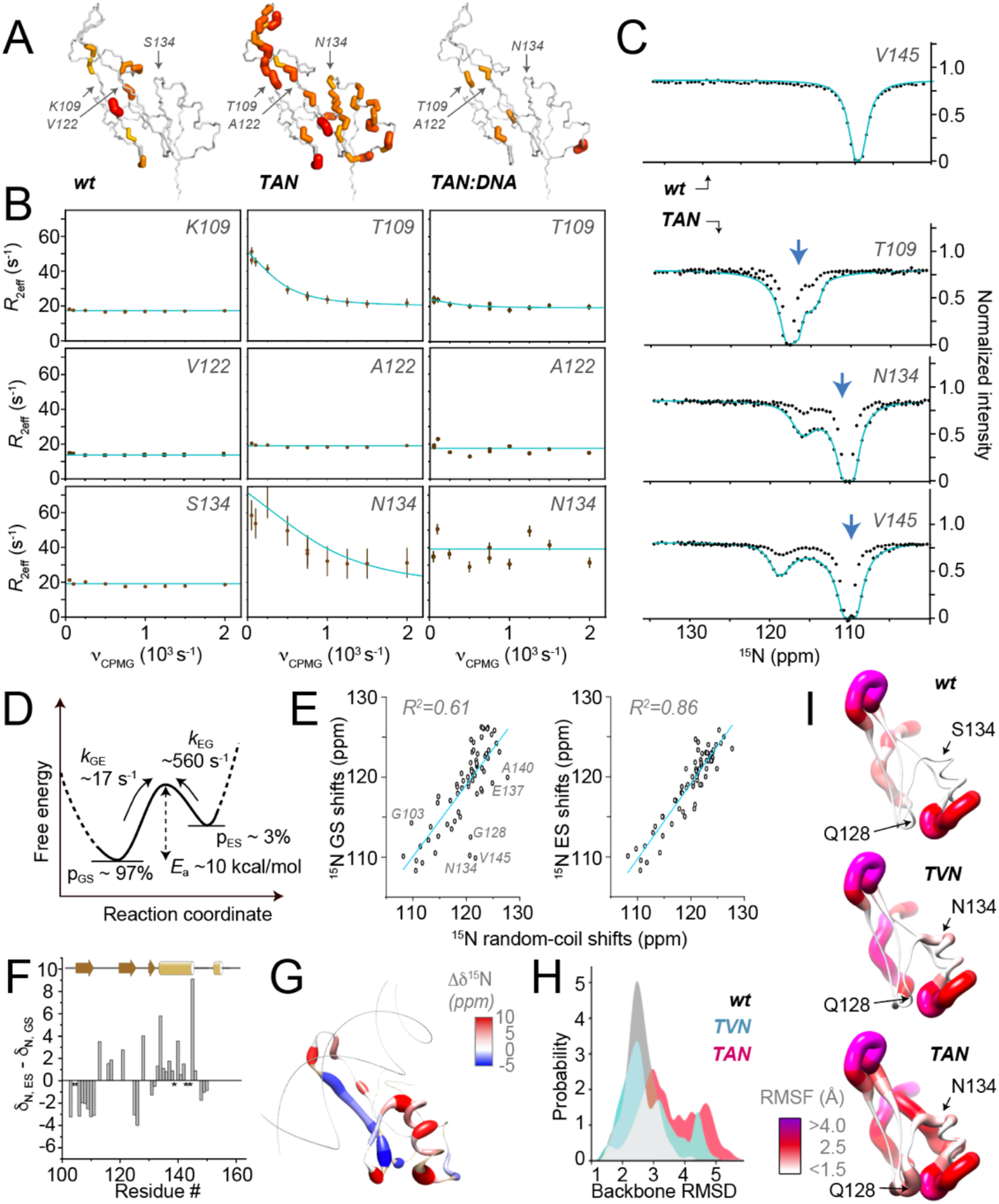
Conformational destabilization of the hmC/mC reader. **A)** Comparison of exchange contributions *R*ex from CPMG RD experiments for apo wt, apo TAN, and TAN:DNA complex, mapped onto the MBD structure (pdb 3c2i). **B)** Constant-time ^15^N CPMG profiles of the wt mC/mC reader (left), the hmC/mC reader TAN (middle), and the TAN:DNA complex (right) for the three mutation sites at 18 °C. **C)** ^15^N CEST profiles (at 10 °C) of TAN, recorded with a *B*_1_ saturating field of 15 Hz (no fit) and 30 Hz (cyan fit) of these residues, in comparison with the wt, taking V145 as a representative residue (top). The blue arrows denote ^15^N shifts of the protein in the DNA-bound state (at slightly higher temperature of 32 °C, see more details below). **D)** Thermodynamics and kinetics of conformational exchange as derived from CEST data and their temperature dependence. **E)** Correlation between either ground-state (left) or excited-state chemical shifts with neighbor-corrected ^15^N random-coil shifts, revealing the tendency of unfolding in the excited state. **F)** and **G)** Difference between ground-state and excited-state ^15^N chemical shifts in apo TAN, plotted as a function of sequence and mapped onto the structure (3c2i), respectively, with consistent trends towards partially unfolding. **H)** Distribution of deviations (RMSD) of the backbone from the X-ray structure, demonstrating a slight destabilization of apo TAN compared to the others in MD simulations. **I)** Root-mean-square fluctuations (RMSF) witnessed in MD simulations of apo proteins, with fluctuations of the regions around residues 134 and 128 “switched on” successively going from wt to TVN to T AN (top to bottom).

To more closely assess the conformational states sampled by the apo hmC/mC reader during the exchange, we used chemical-exchange saturation transfer (CEST).^20^ Here, a weak radiofrequency field is applied to capture chemical shifts of minor conformations for each amide site. We recorded ^15^N-based CEST on TAN at temperatures of 291, 283, and 278 K. At 283 K, we performed CEST analysis at two different saturating fields, 15 and 30 Hz (Fig. 4C). A large dip represents the dominant, lower-energy ground state (GS), whose ^15^N chemical shifts coincide with the regular NMR resonances, and a smaller dip for many residues discloses the chemical shift of an energetically excited state (ES). Amides with a significantly different minor dip, likely involved in strong structural transitions, are again distributed throughout the protein. Fitting the data to a two-state model (see the SI) indicates a dynamic equilibrium, GS ≠ES, on the ~1 ms timescale, with a total exchange rate *kex* of 572 s^-1^ at 283 K and an excited-state population *p*_ES_ of around 3% (Fig. 4D). ^15^N CEST also allowed to derive forward and backward rates, *k*_GE_ and *k*_EG_, of 17 s^-1^ and 555 s^-1^, respectively (SI). This is in congruency with global fitting of the CPMG data at 283 K for TAN, which yielded a similar *k*_ex_ of 458 ±68 s^-1^. Further ^15^N CEST profiles of TAN are shown in Fig. S9. An Arrhenius plot of *k*_ex_, derived from CEST at three temperatures (291, 283, and 278 K, Fig. S10), provides an estimate for the activation energy of 9.5 kcal/mol, which is in line with the timescale of exchange observed in the RD data. Many residues with substantial discrepancies between ground- and excited-state chemical shifts are located in or near the binding-loop and a1 and denote changes in particular for residues just before and after the first helix (Figs. 4E-G). This agrees with the contradictory RDCs for this helix and hence ambiguous relative orientation of the helical residues with respect to the β-sheet.

Noteworthily, the absolute change in ^15^N shifts for the excited state tends to be downfield for helical residues, in particular the beginning (N134) and end (V145) of a1, and upfield for β1, consistent with a temporary, partial release of the secondary structure at these sides (Figs. 4F/G). Accordingly, the excited state represents a partly molten conformation, whose temporary adoption becomes possible due to altered interactions of the hydrophobic side chain in position 122 in the interface between the long β-sheet, α-helix, and C-terminal residues.

On the other hand, in line with its temperature-dependent HSCQ spectra, the double mutant TVN does not show the strong conformational exchange in ^15^N CEST or CPMG experiments (Figs. S11 and S12, respectively). The onset of vast chemical exchange by (and only by) V122A in the (otherwise identical) triple mutant protein supports the notion that the V122A mutation allows to modulate the conformational-exchange dynamics, which – when incorporated in addition to the constitutive changes in the DNA binding interface (K109T, S134N) – ultimately enables the reader to achieve high-affinity binding to the hmC/mC.

In order to shed further light on the structural fluctuations with atomic resolution, we interrogated dynamics in the apo proteins (wt, TVN, and TAN) and their DNA complexes (see section on DNA binding below) in MD simulations, which provided a detailed description of motion up to 2.5 μs. Even though slower motions on the μs timescale and beyond cannot be faithfully sampled (unless enhanced-sampling techniques are used, which render the interpretation of timescales challenging^21^), the tendencies seen in the MD simulations qualitatively match those observed experimentally. Figs. 4H and I show the distribution of RMSDs to the X-ray structure (over all residues) and the residue-resolved root-mean-square fluctuations (RMSFs) of the apo proteins, respectively. Whereas the region around residues 132 and 138 shows increased plasticity over wt for both TVN and TAN mutant, TAN shows an additional systematic increase of backbone fluctuations between 119 and 132. A similar increase is observed between 103 and 109, preceding mutation K109T. Interestingly, we do observe the above-described local unfolding seen experimentally also in one of our five MD trajectories of TAN. Even though this is only a single (but reversible) event (due to the limited MD timescale), and hence has to be interpreted very carefully, it may shed further light on the destabilization of the structure by V122A mutation. Fig. S13 shows an overlay of the structures, where β1/β2 (harboring T109) is slightly reoriented relative to the rest of the structure (starting from Y132), L1 loop becomes extended, and a1 is shortened by one turn (reaching only up to G141). Extended simulations, combined with enhanced sampling methods, will be required to further characterize the nature of the putative locally unfolded state.

### The structure of the hmC/mC reader in complex with DNA

In addition to the apo proteins, the complex between TAN and hmC/mC DNA was subjected to NMR investigation of structure and dynamics. To more closely investigate the binding of the hmC/mC reader to its DNA, ^15^N-labeled TAN was titrated and equilibrated with an unlabeled, double-stranded hairpin oligonucleotide, carrying asymmetric hmC/mC modifications in the central CpG dyad, in a 1:1 molar ratio. We completed sequence-specific backbone and sidechain assignments of the TAN-DNA complex by similar experimental strategies as for the apo proteins (Fig. S14, deposited under BMRB accession code 34745). We then assessed chemical-shift perturbations (CSPs, see the SI for details) upon complex formation, which are shown as a function of sequence and depicted on the X-ray structure of the wt reader (pdb 3c2i) in Figs. 5A and B, respectively. CSPs largely match the positions expected from the wt complex (pdb 3c2i), with perturbations seen in particular for the poorly structured loop L1, which slides into the DNA major groove. The ^1^H^ϵ^ of R111 shows its ^1^H resonance strongly downfield-shifted from 8 ppm to 9.3 ppm. Similarly, G114 ^15^N is moved upfield from 109.2 ppm to 102.6 ppm, A117 ^1^H^N^ from 7.36 to 6.84 ppm, R111 ^15^N^η^ is shifted downfield by 11 ppm, and N134 H^δ^ is shifted by around 0.3 ppm (Fig. S15). The tertiary structure of the complex was elucidated using a similar set of restraints as for the apo structure; however, RDCs for the complex could not be obtained due to sample instability in the presence of alignment media. Also, due to the absence of intermolecular NOEs, we did not specifically include the DNA in structure determination. Within the precision of this assessment, all individual structural elements of the TAN:hmC/mC complex, deposited as PDB 8ALQ, seem highly reminiscent of the wt reader in complex with a symmetrical mC/mC dyad as observed in 3c2i (see an overlay of lowest-energy NMR conformers in Fig. 5C).

**Fig. 5:**
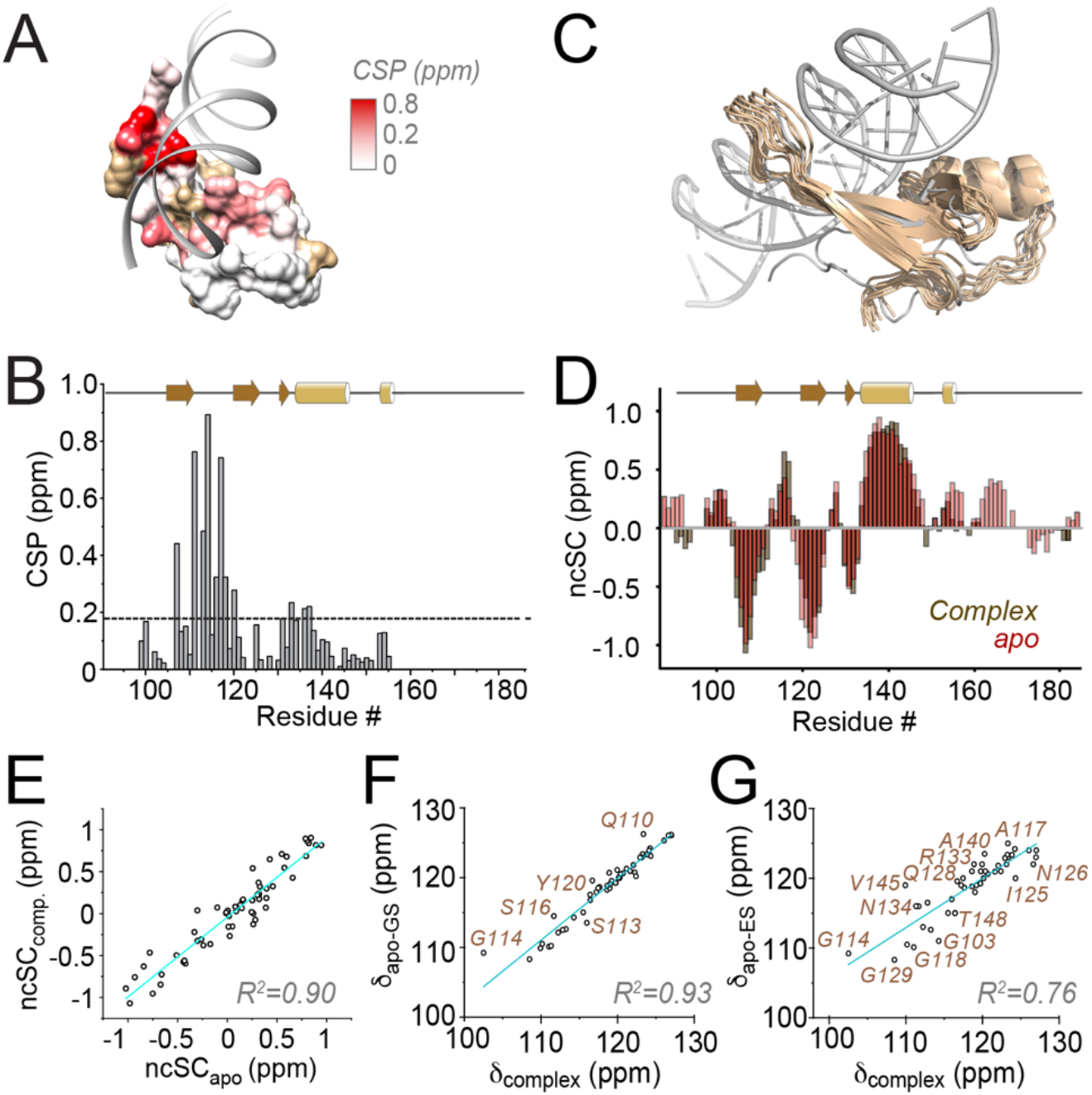
Experimental observations for structure and interactions in TAN complexed with its target hmC/mC DNA. **A)** and **B)** Chemical-shift perturbations (combined ^1^H and ^15^N shifts, see the SI for details) in the hmC/mC reader upon complex formation displayed on 3c2i and as a function of sequence, respectively. Shades of red denote CSPs above 0.2 ppm, golden patches missing assignments in either apo protein or complex. **C)** NMR structure of the TAN:hmC/mC complex, displayed as a bundle of the 10 lowest-energy conformations, overlaid with 3c2i (gray). **D)** Overlay of neighbor-corrected secondary-structural propensities, derived from C^α^, C^β^, CO, N, and H^α^ shifts and their random-coil values, of TAN in apo form and in complex with hmC/mC DNA. **E)** Correlation of secondary-structural propensities with those of the apo protein. **F and G)** Correlation of ^15^N shifts in the complex with either ground-state or excited-state shifts in apo-TAN, respectively.

We asked whether the bound form resembled either the compact or the destabilized excited state of the TAN conformational ensemble. All secondary structural features of the complex are essentially identical to the ground state apo protein (Figs. 5D and E). More interestingly, knowing the residue-resolved ^15^N chemical shifts of the protein in the complex, correlations were sought between these and ^15^N shifts of either the ground or excited state obtained from CEST data of apo TAN (Figs. 5F and G, respectively). Again, a correlation with a correlation coefficient *R*^2^ of around 0.93 for the ground state shifts, in contrast to much larger deviations to the partially unfolded state (*R*^2^ of 0.76), shows the ground state secondary structure of the apo reader to be reconstituted upon binding. This speaks against conformational preselection via a defined excited state as the main mechanism and instead points to facilitation of induced-fit binding of the apo protein to its target by decreased rigidity. Associated with a high energy barrier, this plasticity, however, incurs low entropic costs upon complex formation.

### Increased plasticity allows facilitated DNA binding

We closely inspected the MD simulations of the complex to complement the experimental data on structural properties, dynamics, and interactions and further elucidate the specificities of the interaction of TAN as the first hmC/mC reader with its target DNA on the atomic level. As an expected source of adopted selectivity in addition to the known interactions of the wt protein,^22^ the S134N mutation is confirmed to allow specific H-bonding between 5-hmC and reader (Figs. 6A and S16). (The SI also contains a short movie depicting this interaction.) H-bonding both relates to the hmC hydroxyl group and the phosphate backbone (Fig. 6B). In addition, we again inspected how complex formation is facilitated by specific structural features of the TAN mutant. Indeed, the formation of the TAN:hmC/mC DNA complex in the simulation is associated with slight conformational rearrangements. Whereas some rearrangement also occurs upon wt:mC/mC complex formation, the specific characteristics differ. Fig. 6C visualizes the nature of displacement via an overlay of the average apo structure and the respective MBD:DNA complex (aligned with respect to the β1/β2 sheet). α1 in the average structure of the TAN:hmC/mC complex has slightly reduced absolute displacement (1.9 Å) from its position in apo TAN, both in comparison to the wt:mC/mC complex (2.8 Å) as well as the TVN:hmC/mC complex (3.3 Å; displacement of the helix, determined using F142 CO coordinates). However, with the slight differences adopted in its apo tertiary structure (Fig. 3E and S17B), the displacement needed in TAN is a *parallel slide* of the helix, whereas for the wt, a mere *bending* is necessary. (TVN shows an intermediate behavior, where both is necessary to accommodate hmC/mC DNA.) With its decreased rigidity of the core and hence reduced strains for the architecture of its new binding interface, TAN – upon sliding into the major groove of the DNA – is able to adopt the exact structural characteristics required for the above interactions with the hmC/mC dyad to form. MD pinpoints this elevated plasticity (Fig. 6D, left) to β1 (102-110, allowing to tune R111 interactions with guanosine) as well as the inter-subunit interface (118-140, which includes K130 interactions to the phosphate backbone, R133 interactions with guanosine, and N134 H-bonding to hmC), in complete agreement with the dynamics from the NMR experiments described above. This plasticity facilitates stable and tight interactions with hmC/mC DNA, upon which this mobility ceases (Fig. 6D, right). Note that in contrast to the now stably formed interactions of the TAN complex, for the more restrained TVN double mutant, increased local fluctuations are now observed at residues 126-130, denoting a weakened K130-phosphate interaction, when complexing hmC/mC DNA (Fig. 6D, dark blue curve, and S17A, right panel).

**Fig. 6.:**
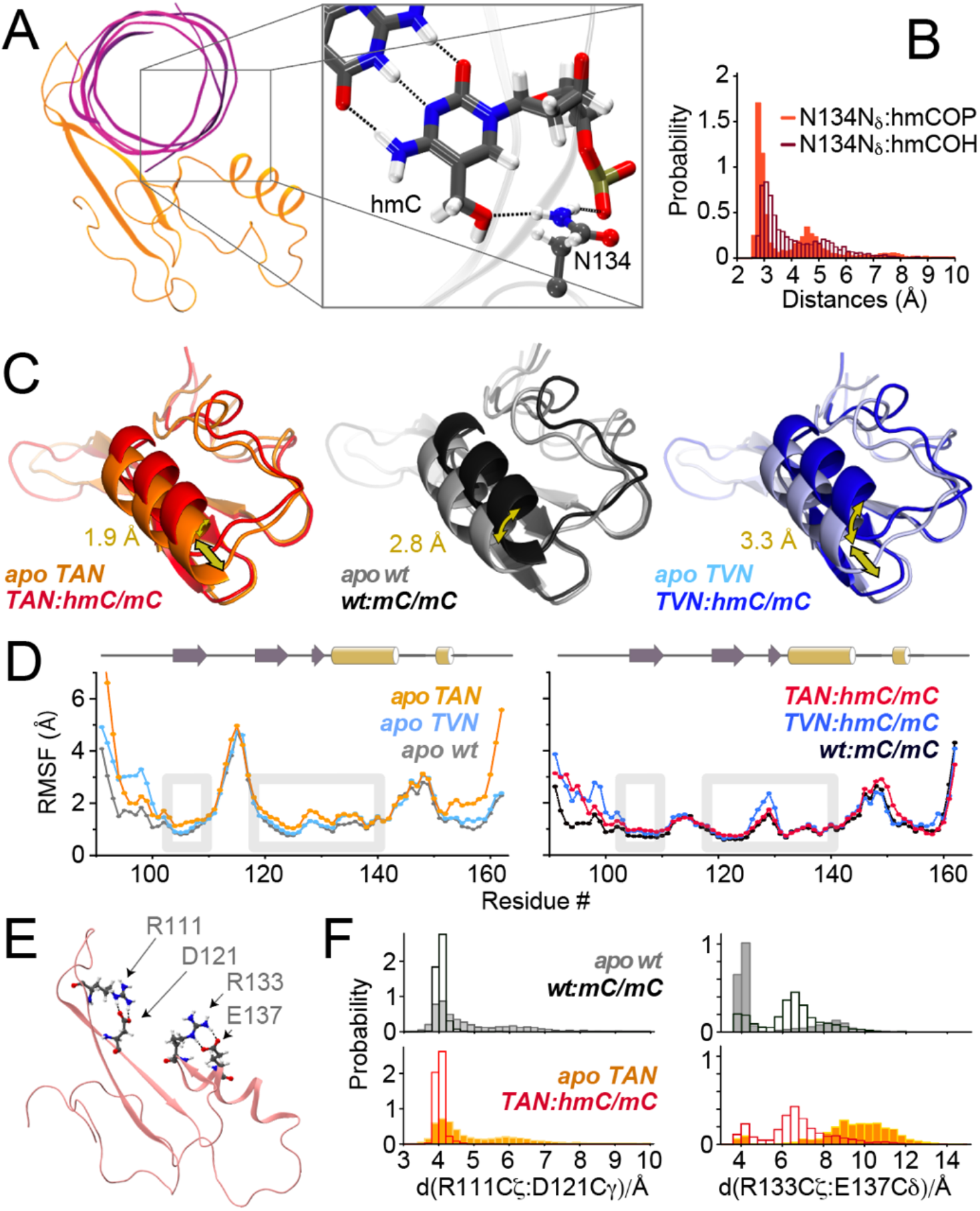
In-silico observations regarding complex formation between MBD and target DNA for different constructs. **A)** Formation of the N134:hmC H-bond as the enthalpic driving force in case of the epigenetically modified CpG. **B)** Histograms of N134 Ns H-bonding to hmC hydroxyl and phosphate. **C)** Average backbone structures from 2.5 μs MD simulations of wt, TAN, and TVN in apo form or in complex with their target DNAs, aligned with respect to β1/β2. The annotated arrows reflect the displacement between the F142 carbonyl (CO) group in apo proteins and complexes. **D)** Residue-specific RMSFs in apo proteins and complexes. Gray boxes highlight β1, framing the important R111:guanosine interaction, and the area involving the inter-subunit interface (res. 122: central mutation site, 130: Lys to phosphate backbone, 133: Arg to guanosine, 134: H-bond to hmC, 137: potential R133 preorganization). **E)** Arg residues forming a main selectivity aspect, including differentially preorganizing salt bridges. **F)** Distance probabilities of the R111/D121 (left) and the R133/E137 salt bridge (right) of wt (top) and TAN (bottom) in apo form (solid bars) and in complex with matching target DNAs (open bars).

Two significant interactions by Arg “fingers” 111 and 133, which can form H-bonds with the DNA backbone and hence constitute a major enthalpic driving force for complex formation, have been discussed (Figs. 1C, 6E and S16).^14^ These H-bonds are stably formed in our simulations (Fig. S18). The large perturbation of R111 chemical shifts upon binding (see above) increases the discrepancy that its ^1^H^Ϗ^ proton has in the apo state compared with other Arg sidechain moieties (Fig. S19A). This deviation, both for wt and TAN, confirms a preorganization of the salt bridge with neighboring D121 in the apo state that eventually also characterizes the complex of the wt mC/mC reader with DNA in crystals. Fig. S19B shows strips of a ^15^N-edited NOESY experiment, where the preformed intraresidual contacts between R111 and D121 in the TAN mutant apo form are apparent. MD data for R111, which maintains a stable salt bridge with D121 both in apo and complex states of *all* readers in the simulations, is shown in Fig. 6F, left panels. Importantly, in contrast to R111 and to what has been proposed in the framework of a possible selectivity mechanism,^6^ the salt bridge preorganizing Arg133 for its H-bond to guanosine in the complex is witnessed in the apo wt protein, but it is released in our simulations of the wt:mC/mC complex (Fig. 6F, upper right panel). In the TAN mutant, by contrast, the R133/E137 salt bridge is absent both in the apo form and in the complex (Fig. 6F, lower right panel, and Fig. S20). Absence of this intramolecular ionic lock avoids energetic penalties to open it upon complex formation, apparently without corrupting the formation of the important R133 H-bond. Finally, the precise positioning of R111 hinges on strand β1. Flexibilization of the relative position of this strand (Figs. 4B and 6D left) by Lys-to-Thr mutation of residue 109 may reduce strains negatively associated with R111 binding in the context of the modified interface.

Overall, a high affinity of the new MBD to the asymmetric, hydroxymethylated dyad seems to depend on smoothening of the free-energy landscape of the reader towards enabling conformational adaptations. This key property is fine-tuned by interactions between the three central protein secondary-structural elements (helix α1, the extended β-sheet, and the domain connecting the two) – which is defined by the central residues in the hydrophobic core – rather than the binding interface itself.

## Discussion

The above results show that redirecting the specificity of MBDs as a naturally existing scaffold to a new epigenetic CpG duplex modification hinges on tailored modulation of the thermodynamics of binding, derived not only from new intermolecular contacts but also tuning the characteristics of conformational plasticity in the interface. The enabling plasticity, adjusted via central residues within the protein core, derives from changes in the steric matches in the inter-domain interaction surfaces. The extent of this mobility, apparent from a tendency towards rare local unfolding, is strongly enhanced in the new hmC/mC reader. The adoption of an excited state on the μs-ms timescale motion and hence the presence of a similar tendency, albeit less pronounced, also characterizes the natural mC/mC reader. Suitably adjusted plasticity of the binding interface thus seems an important general aspect of target recognition for the MBD fold. At first glance, such “disorder” in the apo scaffold seems like an entropically disadvantageous property for a high-affinity binder in which plasticity decreases upon binding. However, with features of this plasticity being very modestly tuned (i. e., with maintained, well-defined secondary-structural elements, tight conformational restrictions, and a high activation barrier), the penalty at physiological temperatures is minor – while still redefining a 25 Å wide binding interface – and can be largely compensated by maximized enthalpic gains due to optimized H-bond formation.

The design of reader proteins that can serve as probes for the analysis of postsynthetic modifications of nucleic acids constitutes a current key aim of the soaring fields of epigenetics and epitranscriptomics. We showed that sought new properties of relevant protein-nucleic acid interfaces can be induced by directed evolution based on natural progenitors. Whereas the design and selection of mutant libraries with well-defined randomization sites guided by visual inspection of crystallographic structures that report on local interactions is the most intuitive approach, our data show that, by contrast, interrogating nucleic acids by designed reader proteins can also critically hinge on correctly adjusting protein plasticity as a modulator of selective complex formation. As such, central mutation sites far from interaction surfaces but relevant for interdomain connectivity allosterically can enable high affinity and selectivity of readers, hence allowing for unexpected new perspectives for progress, particularly in the new field of CpG duplex mark recognition. We believe that the interrogation and understanding of dynamic networks in epigenetic readers, writers, and erasers can represent a fundamental element for designing future probes to decipher and effectively modulate the layer of epigenetic control of cell fate and function. These findings will be interesting for design problems in other contexts of target recognition, as – despite active research towards understanding of dynamic networks^23–25^ – dedicated allosteric optimization of large-scale dynamics as a lever for any desired functionality still tends to escape awareness in the creation of new molecular tools.

## Conclusion

Here we have demonstrated that the high affinity and selectivity of the first designed epigenetic reader for oxidized CpG dyads is leveraged by well-defined conformational plasticity of the DNA binding interface, remotely orchestrated by interactions in the hydrophobic core. The revelation of an intermediate-timescale conformational exchange towards a partial melting of secondary-structural strains, elucidated via extensive NMR and MD interrogation of protein structure and dynamics, demonstrates adapted plasticity as a lever for specific reader:DNA interactions within the vast landscape of differential epigenetic modifications. Albeit to a lower extent, exchange dynamics on the same timescale are also visible for the natural, mC/mC-specific MBD. Our study suggests that the aspect of tailored conformational plasticity may both, help understanding physiological reader selectivities and facilitate the design of novel readers as specific molecular probes for different CpG duplex marks. The revelations will propel the advances in the emerging field of DNA recognition and thus in deciphering the elusive roles of individual CpG duplex modifications in chromatin regulation.

## Acknowledgements

Funded by the Deutsche Forschungsgemeinschaft (DFG, German Research Foundation) – 27112786, 325871075, and the Emmy Noether program. Funded by the Deutsche Forschungsgemeinschaft (DFG, German Research Foundation) under Germany’s Excellence Strategy - EXC 2033 – 390677874 – RESOLV, and EXC-114 – 24286268 – CiPS-M. Funded by the European Research Council (ERC CoG EPICODE, No. 723863 to D.S.)

